# *miR-124* acts during *Drosophila* development to determine the phase of adult circadian behavior

**DOI:** 10.1101/2025.02.07.637081

**Authors:** Yongliang Xia, Chenghao Chen, Patrick Emery

## Abstract

Circadian behaviors need to be properly phased with the day/night cycle to be beneficial. We previously showed that the microRNA *miR-124* regulates circadian behavior phase in *Drosophila*. Here, we report that *miR-124* expression during larval development is required for proper phasing of both morning and evening peaks of activity in adults. Loss of *miR-124* results in significant miswiring within the circadian neural network and severely alters neural activity rhythms in the ventral Lateral Neurons (s-LNvs) and the posterior Dorsal Neurons 1 (DN1ps), which control the timing of morning and evening activity. Silencing the s-LNvs in *miR-124* mutant flies restores the phase of evening activity, while activating the DN1ps rescues the phases of both morning and evening activities. Our findings thus reveal the pivotal role of *miR-124* in sculpting the circadian neural network during development and its long-lasting impact on circuit activity and adult circadian behavior.

## Introduction

Most living organisms, including bacteria, fungi, plants, insects, mammals, and humans, experience daily changes in their surrounding environment, such as variations in light and temperature^1,2^. To adapt to and anticipate these environmental cycles, they have evolved an internal timekeeping system known as the circadian clock. This system helps organisms orchestrate their temporal relationship with the ever-changing environment, enhancing their efficiency in resource utilization and activity scheduling. However, circadian clocks remain rhythmic with a period of ca. 24-hour even under constant environmental conditions, such as constant darkness (DD). Thus, this intrinsic rhythm allows organisms to sustain biological processes that rely on timing, ensuring continuity in their behavioral patterns and physiological responses, even in the absence of external Zeitgebers^3–5^. These circadian clocks consist of self-sustained transcriptional feedback loops that function as endogenous oscillators, driving the rhythmic expression of a wide range of genes involved in diverse biological processes, from metabolic pathways to physiology and complex behaviors^6,7^. In *Drosophila melanogaster*, the molecular clock comprises three interconnected feedback loops^8^. In the core loop, the transcriptional activators CLOCK (CLK) and CYCLE (CYC) dimerize via PAS domains to drive the transcription of two genes encoding transcriptional repressors, PERIOD (PER) and TIMELESS (TIM). These repressors form a complex that translocates into the nucleus and represses CLK/CYC activity^9,10^. This molecular clockwork is found in approximately 240 neurons in the adult *Drosophila* brain, which can be classified into several subsets based on their anatomical localization and size^11,12^. Different groups of clock neurons display distinct gene expression patterns^13^ and phases of neural activity, which regulate specific aspects of daily locomotor rhythms, such as the timing of morning and evening activities^14–16^. Through communication via neuropeptides and neurotransmitters, the ca. 240 clock neurons form a plastic network that adapts to different environmental conditions^17,18^.

The synchrony between intrinsic circadian clock and rhythmic environmental cues provides an adaptive advantage, while their misalignment has detrimental effects on survival and health, including in humans^19–21^. Evidence from various species also shows that mutants with altered circadian periodicity, when placed in non-matching environmental cycles, experience a reduction in lifespan. In contrast, synchrony between the circadian clock and environmental cycles extends lifespan and improves fitness^3,22–24^. Both in nature and in laboratory conditions, individuals with circadian periods closely matching environmental cycles are favored under competition^25,26^. For human beings, besides circadian gene mutations, modern lifestyles significantly contribute to the disharmony between our body clocks and external time. Factors such as untimely eating, shift work, travel across different time zones, social jetlag, and extended artificial lighting disrupt the proper entrainment of our circadian systems to local environmental cycles^20,27,28^. This irregularity prevents our circadian clocks from functioning optimally, leading to various health issues. It is therefore essential to understand how the phase of various circadian rhythms is matched with the day/night cycle.

We now have a good understanding of the mechanisms underlying circadian molecular clocks and their entrainment to environmental cycles. However, the mechanisms by which these clocks generate properly phased circadian outputs, such as behavioral rhythms, is not well understood. Genetic screens conducted for example in *Drosophila* have been remarkably efficient at identifying circadian pacemaker genes. A great number of period altering mutants have been isolated. As expected, such mutants usually do not just affect the period of circadian rhythms under constant conditions, but also their phase when mutant animals are exposed to environmental cycles. Most famously perhaps, flies with long or short period mutations in the *per* gene display delayed and advanced circadian phases under Light:Dark (LD) cycles, respectively^29,30^. Gene mutants that exclusively affect the phase of a circadian rhythm, without impacting the period of the underlying circadian pacemaker, are rather rare. This explains why our knowledge of the mechanisms by which the circadian pacemaker determines the phase of overt circadian rhythms is quite limited.

Circadian behavior in *Drosophila* is characterized by two peaks of activity at dawn and dusk. These peaks are observed under both LD and DD conditions, although the morning peak of activity tends to progressively fade under constant darkness^31^. We and others previously reported that the loss of *miR-124* in *Drosophila* leads to an advanced evening phase under DD conditions, but importantly does not impact the period and phase of the molecular clock in the circadian neurons that controls rhythmic behavior^32,33^. *miR-124* thus provides an entry point to elucidate the mechanism of circadian phase determination. In this study, we utilized CRISPR/Cas9 to disrupt *miR-124* in neurons. This approach revealed that *miR-124* functions during development to regulate circadian phase. Additionally, the loss of *miR-124* resulted in important miswiring of the circadian circuits, and abnormal activity of neurons controlling morning and evening activities. These findings suggest a critical role for *miR-124* in the development and proper functioning of the circadian neural network.

## Results

### Loss of *miR-124* causes opposite shifts in morning and evening phases

Previous studies showed that loss of *miR-124* results in an advanced phase of evening activity under DD, without affecting the free-running period of activity rhythms^32,33^. However, under a 12 h/12h LD cycle, *miR-124^KO^*flies display a completely normal evening peak phase, indicating that light input can correct the phase defect. This likely occurs through plastic recruitment of different evening neurons under LD and DD^18^.

To further investigate the effects of light on circadian phase, we examined the circadian locomotor activity of *miR-124^KO^* flies under different LD cycles. Consistent with previous studies^32,33^, under a regular LD cycle (LD12:12), the phase of evening activity was not affected in *miR-124^KO^* flies, although its amplitude was reduced. Also, both morning activity and the startle response resulting from the light-on transition were absent in the mutants (Fig. 1A and 1A’). Nocturnal activity was also elevated in these mutants. Following release into DD, *miR-124* mutants displayed as previously observed^32,33^ a unimodal activity pattern with a markedly broad and ca. 5h advanced evening peak (Fig. 1B and 1B’). These defects were corrected with a transgene containing *miR-124*. Similar phenotypes were observed under long photoperiods (LD16:8). Both the morning anticipation and the light-on startle response were lost, but the evening activity phase was comparable to that of wild-type flies (Fig. S1). In contrast, *miR-124^KO^*flies showed robust morning activity under short photoperiods (LD 6:18).

**Fig. 1.**
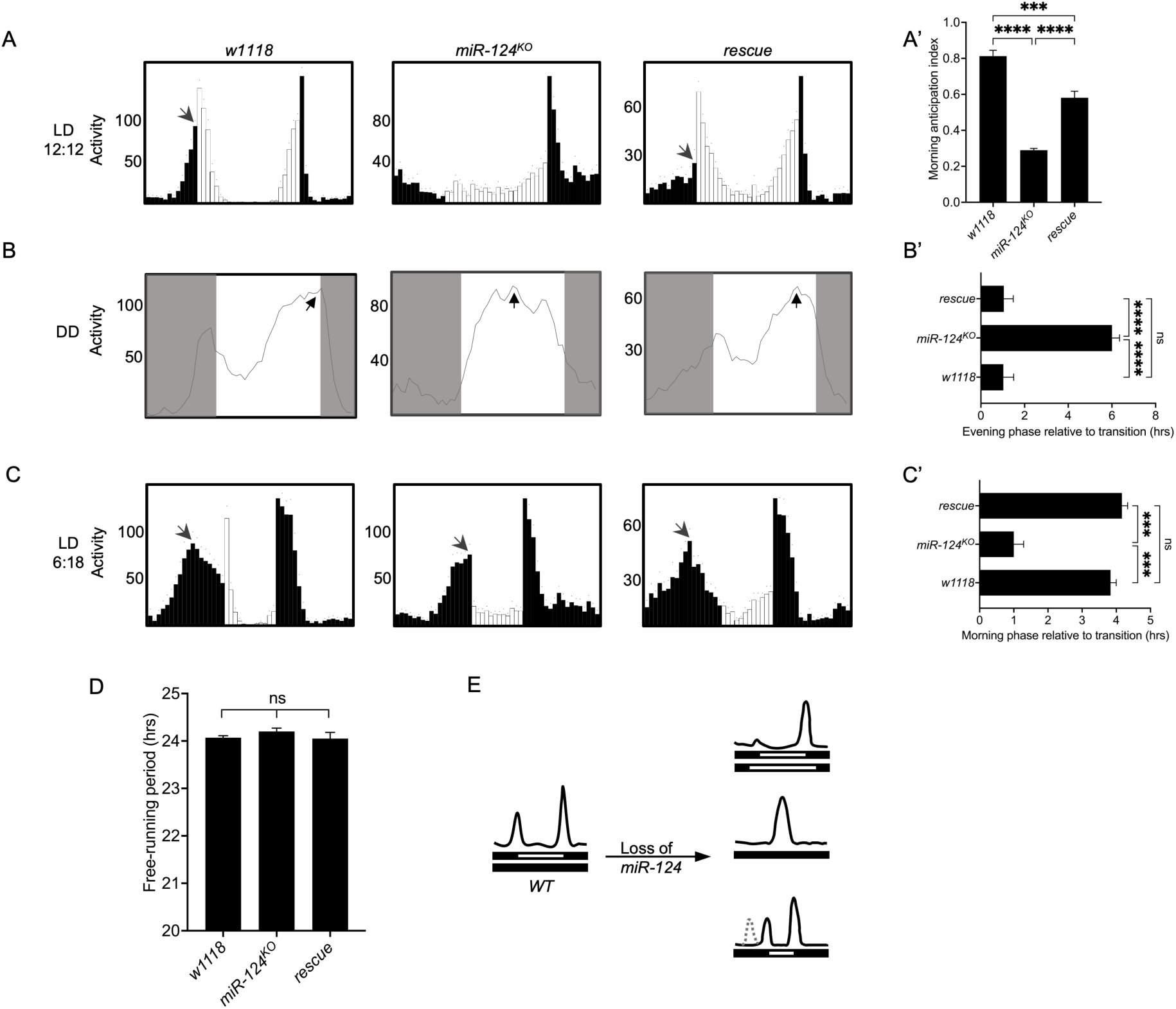
Loss of *miR-124* shifts morning and evening peak phases in opposite directions. *A-C*, Panels depict behavior of *w*^1118^, *miR-124*^KO^, and rescued *miR-124*^KO^ male flies. The rescued flies contain a 19 kb genomic transgene [*39N16*]. Open arrows indicate the morning peaks of activity, and regular arrows the evening peak of activity. This will be the case throughout this report. *A*, Averaged locomotor activity profiles under 12:12 LD cycle. White bars indicate activity during day, black bars at night (n ≥ 10 per genotype). *A’*, Quantification of morning anticipation index under 12:12 LD. *B*, Averaged locomotor activity profiles under DD (n ≥ 8 per genotype). Gray shades represent the subjective night. *B*, Quantification of evening peak phase under DD, relative to the subjective light-off transition. *C*, Averaged locomotor activity profiles under short photoperiods (6:18 LD) (n ≥ 10 per genotype)*. C*’, Quantification of morning peak phase under short photoperiods is plotted, relative to the light-on transition. *D*, Quantification of free-running periods over five DD cycles. *E*, Schematic diagram summarizing the distinct locomotor activity patterns of *miR-124*^KO^ mutants compared to WT flies under different environmental conditions. Related supplemental data can be found in Figure S1. White bars indicate light phase (day), black bars indicate dark phase (night). Dotted line indicates morning activity peak of WT flies. Quantifications are mean ± SEM from at least 3 independent behavioral experiments, and behavioral panels are from a single representative experiment. This will be the case throughout this report. One-way ANOVA with Tukey’s post hoc test. ***, P<0.0002; ****, P<0.0001; ns, not significant.

However, its phase was severely delayed compared to wild-type controls and rescued *miR-124* mutant flies (Fig. 1C and 1C’). Thus, while loss of *miR-124* advances evening activity in DD, it delays the morning peak of activity under LD cycles. We note that a truncated, delayed morning peak of activity was actually also detectable in *per^S^;miR-124^KO^* double mutant flies in our previous study^33^. We thus conclude that *miR-124* mutants can display a delayed morning peak of activity, as long as its onset falls within the dark period. Under LD12:12, it falls during the light period and is thus suppressed.

In summary, *miR-124^KO^* flies display specific behavior phenotypes under various environmental conditions, due to changes in activity phase. Loss of *miR-124* advances the evening peak in DD, but delays morning activity (Fig. 1E). Because neither the free-running period nor the phase of the neural molecular clocks are affected in *miR-124^KO^* mutants^32,33^ (Fig. 1D), this indicates that *miR-124* specifically regulates the circadian phase of rhythmic behavior.

### Loss of *miR-124* disrupts the activity of circadian pacemaker neurons

Although the molecular clocks of various circadian pacemaker neurons are synchronized, their Ca^2+^ oscillations and thus their neural activity are asynchronous. These Ca^2+^ rhythms usually coincide with locomotor activity bouts mediated by the corresponding neural pacemakers^34^. Since molecular clocks in all pacemaker groups function normally in *miR-124^KO^*flies^32,33^, we investigated whether neuronal activities in circadian clusters were altered in *miR-124^KO^* mutants. We expressed the photoconvertible protein CaMPARI^35^ (Calcium Modulated Photoactivatable Ratiometric Integrator) to monitor the integrated calcium activity of different pacemaker groups over distinct time points. We first detected calcium activity in s-LNvs, as they are responsible for rhythmicity in DD, and dominate under short photoperiods^36^— conditions where *miR-124^KO^* flies show behavioral phase defects. The s-LNvs also control the morning peak of activity, which is phase-shifted in *miR-124* mutant flies. In wild-type flies, Ca^2+^ levels in the s-LNvs peaked near the end of the night, as previously reported, consistent with their role as morning anticipation-driving neurons^15,16,34^. Intriguingly, in *miR-124^KO^* mutants, the Ca^2+^ peak of s-LNvs occurred in the middle of the subjective day under DD (Fig. 2A and 2B), consistent with the advanced evening peak of locomotor activity observed under these conditions (Fig. 1B and 1B’). To determine whether the shifted neuronal activity in s-LNvs is responsible for the phase delay of behavioral activity in *miR-124^KO^* flies under DD, we inhibited the s-LNvs by expressing the inwardly rectifying K^+^ channel Kir2.1^37^ in *miR-124^KO^* mutants. Indeed, repression of s-LNvs in the *miR-124^KO^* was able to correct the evening phase (Fig. 2C and 2D). Note that behavior was observed during the first day of DD only, and we could not assess morning activity phase under LD since inhibition of s-LNvs causes behavioral arrhythmicity under prolonged DD exposure and blunts morning activity^38,39^.

**Fig. 2.**
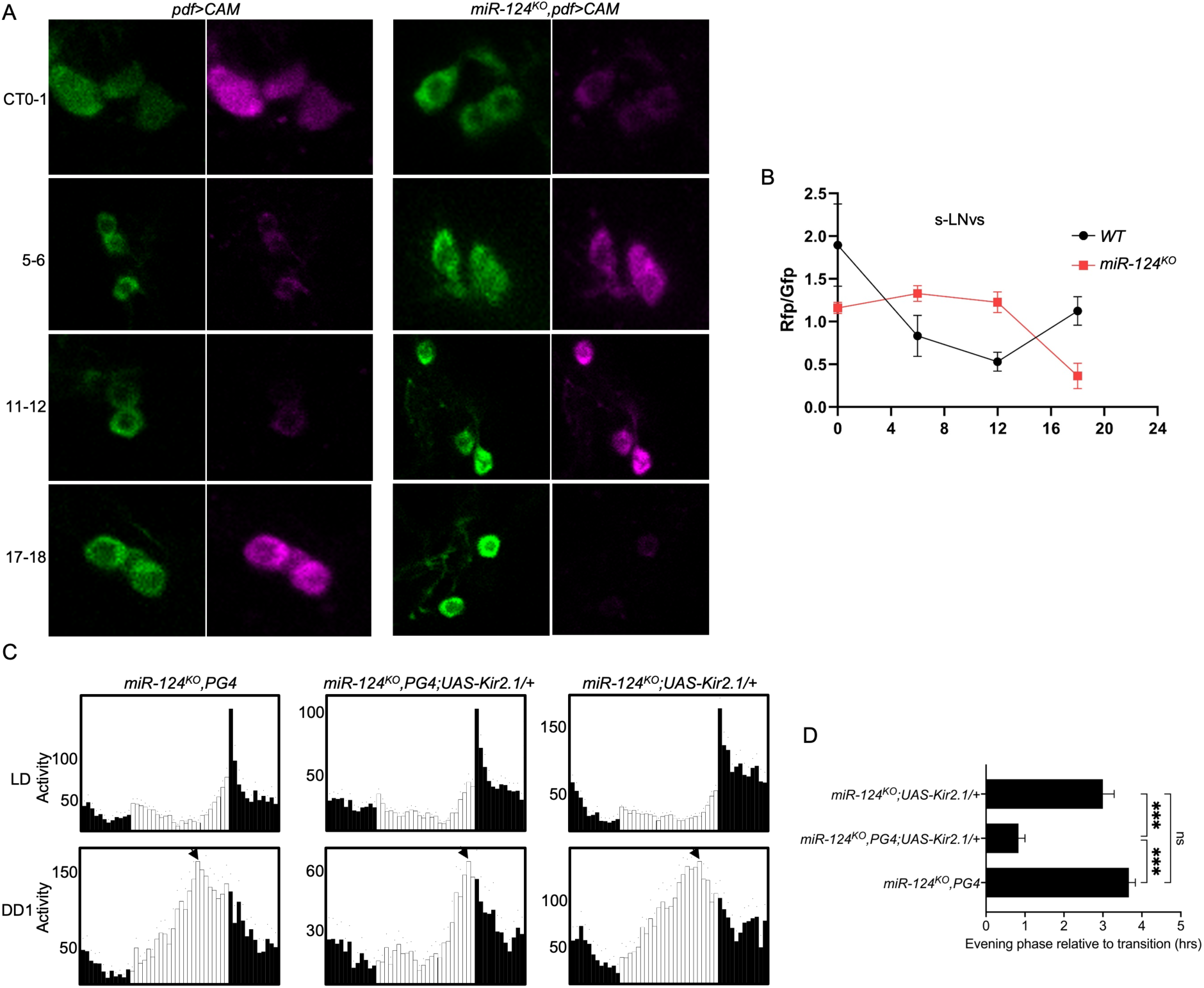
Loss of *miR-124* shifts the neuronal activity of s-LNvs. *A*, Confocal imaging of s-LNv calcium dynamics in WT and mutant brains dissected at different time points (circadian time, CT) during the first day of DD. *B*, Quantification of CaMPARI photoconversion efficiency across the indicated circadian time points (n = 6-8 hemispheres per time point). *C*, Averaged locomotor behavior under three days of LD cycles (upper panel) and first day of DD (lower panel) (n ≥ 10 per genotype). *D*, Quantification of evening peak phase during the first day of DD (N ≥ 3 independent experiments). Error bars indicate SEM in *B* and *D*. One-way ANOVA with Tukey’s post hoc test. ***, P<0.0002; ns, not significant.

A subset of the posterior dorsal neurons 1 (DN1ps), which includes both morning- and evening-activity promoting cells, is directly targeted by the s-LNvs^18,40,41^. We thus also investigated the role of *miR-124* in regulating neural activity in these neurons. In wild-type flies, DN1p-morning neurons (DN1p^M^) showed a Ca^2+^ peak during the late night, while DN1p-evening neurons (DN1p^E^) displayed a Ca^2+^ peak in the mid- to late day. This is consistent with their respective roles in morning and evening anticipatory activities.

Surprisingly, in the *miR-124^KO^* mutants, neural activity was severely depressed at all time points in both DN1p subtypes (Fig. 3A-3D). When we simultaneously activated both DN1p^M^s and DN1p^E^s using the *Clk4.1M-GAL4* driver^40^, the behavioral phenotypes of *miR-124* mutants were partially corrected (Fig. 3E-3G and 3E’-3G’). In LD and DD conditions, *miR-124^KO^* flies now displayed bimodal activity patterns due to the re-emergence of morning anticipatory activity combined with reduction in nighttime activity. Evening and morning activity were of approximately equal amplitude on DD1, and morning activity and evening phase were significantly improved (Fig. 3E’-F’ and 3F’’). Under short photoperiod conditions, *miR-124^KO^*mutants with activated DN1p^M^s and DN1p^E^s showed advanced morning activity phase compared to their controls (Fig. 3G’ and 3G’’). Notably also, *miR-124^KO^* flies, which are as mentioned above more nocturnal than control animals, switched to a wild-type-like diurnal activity pattern upon activation of DN1p neurons. As expected, the lights-on startle responses of *miR-124^KO^* mutants could not be restored by DN1p activation or s-LNv inhibition (Fig. 2C and 3E-3G). Indeed, these responses are not controlled by the circadian clock^42^.

**Fig. 3.**
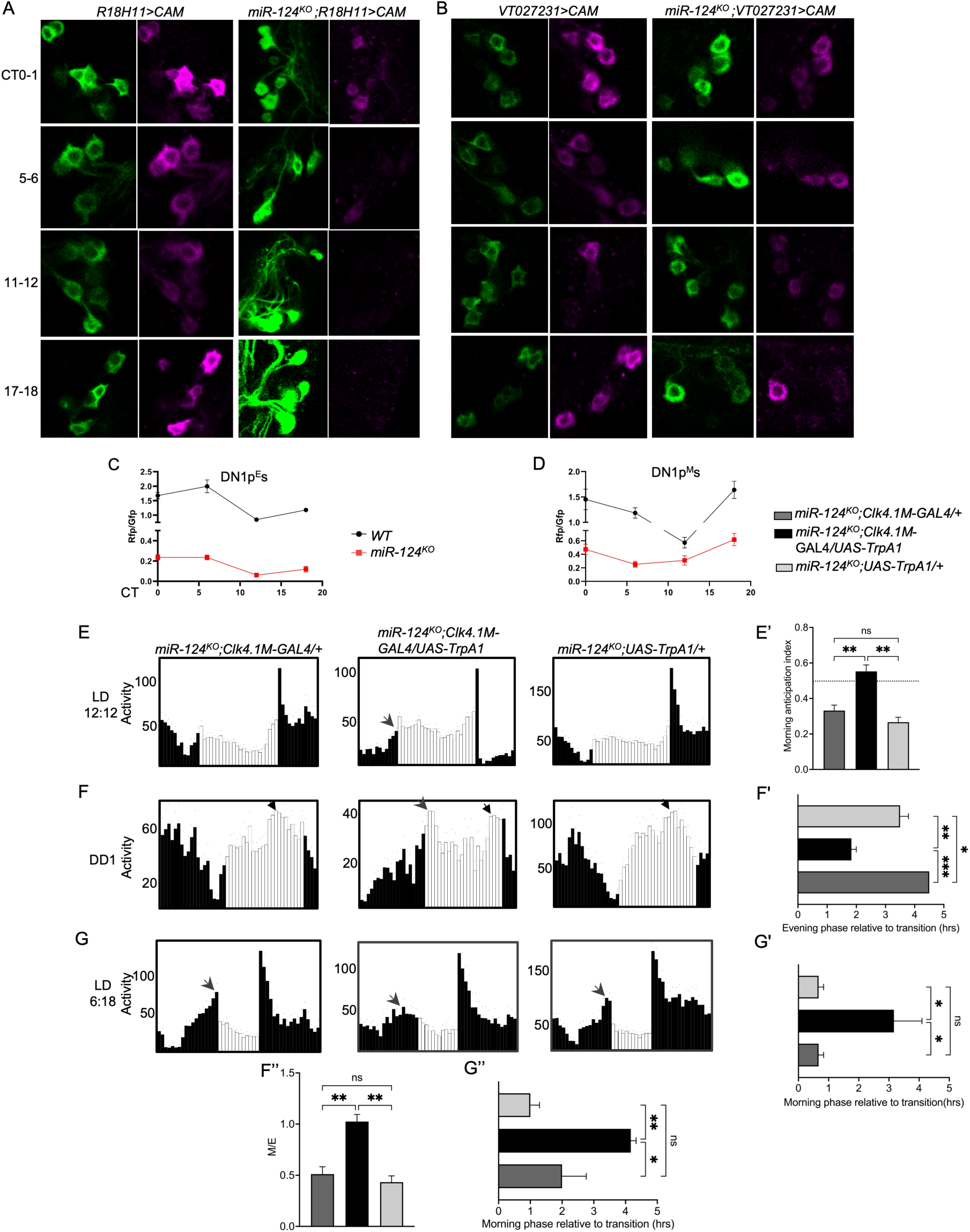
Loss of *miR-124* represses neuronal activity in DN1p^E^s and DN1p^M^s. *A-B*, Confocal imaging of DN1p^E^ (*A*) and DN1p^M^ (*B*) calcium dynamics in WT and mutant brains dissected at different time points (circadian time, CT) during the first day of DD. *C-D*, Quantification of CaMPARI photoconversion efficiency across the indicated circadian time points in DN1p^E^s (*C*) and DN1p^M^s (*D*) (n = 6-8 hemispheres per time point). *E-G*, Representative locomotor activity profiles of flies with activation of DN1ps in 12:12 LD (*E*), first day of DD (*F*) and short photoperiods (*G*) (*E-F*, n ≥ 16 per genotype; *G*, n ≥ 9 per genotype). *E’-G’*, Quantification of morning anticipation index in 12:12LD (*E’*), evening peak phase in DD (relative to the subjective end of the day) (*F’*) and morning peak phase under 6:18 LD (*F’*). *F’’*, Quantification of the ratio between morning and evening peak activity under DD1. *G’’*, Since morning activity was quite flat in *miR-124* mutant flies with re-activated DN1ps, we also measured the time at which 90% of morning peak activity was reached in 6:18 LD. This confirmed that activity phase was indeed advanced in these flies compared to control. Legends for *E’-G’* and *F’’-G’’* are above the panels, next to panel D. Mean ± SEM is shown in *E’-G’*. One-way ANOVA with Tukey’s post hoc test. *, P<0.0332; **, P<0.0021; ***, P<0.0002; ns, not significant. (N ≥ 3 independent behavioral experiments).

Taken together, these results demonstrate that the loss of *miR-124* either shifts or suppresses the neuronal activities of circadian clusters. Silencing s-LNvs or activating DN1ps in *miR-124^KO^* mutants can correct phase defects in morning and evening activities, indicating that *miR-124* regulates circadian phases through these neurons and their activity. Our results also suggest that coordinated antiphasic neural activity in different subset of DN1ps is important for proper phasing of morning and evening activity.

### Pan-neuronal CRISPR/Cas9-mediated disruption of *miR-124* mimics *miR-124^KO^*

To study further the mechanisms by which *miR-124* regulates circadian behavior phase, we turned to conditional CRISPR/Cas9-mediated gene knock-out. Indeed, this approach works efficiently in the *Drosophila* circadian neural network^43,44^. We followed the protocol provided by Port and Bullock^45^, and designed three sgRNAs targeting three different regions of the *miR-124* precursor (*pre-miR-124*) to ensure efficient gene disruption. One of the three gRNAs targeted the coding sequence for the mature *miR-124* (Fig. 4A). Given that *miR-124* is a neuron-specific miRNA^46^, we expressed *miR-124-sgRNAs* and *Cas9* pan-neuronally with the *elav-GAL4* driver. This led to a substantial reduction of *miR-124* level, which were barely detectable in head extracts (Fig. 4B). As expected, *miR-124^CRISPR^* flies showed very similar behavioral phenotypes as *miR-124^KO^* mutants under LD cycles with regular or short photoperiod, and under DD (Fig. 4C-4E).

**Fig. 4.**
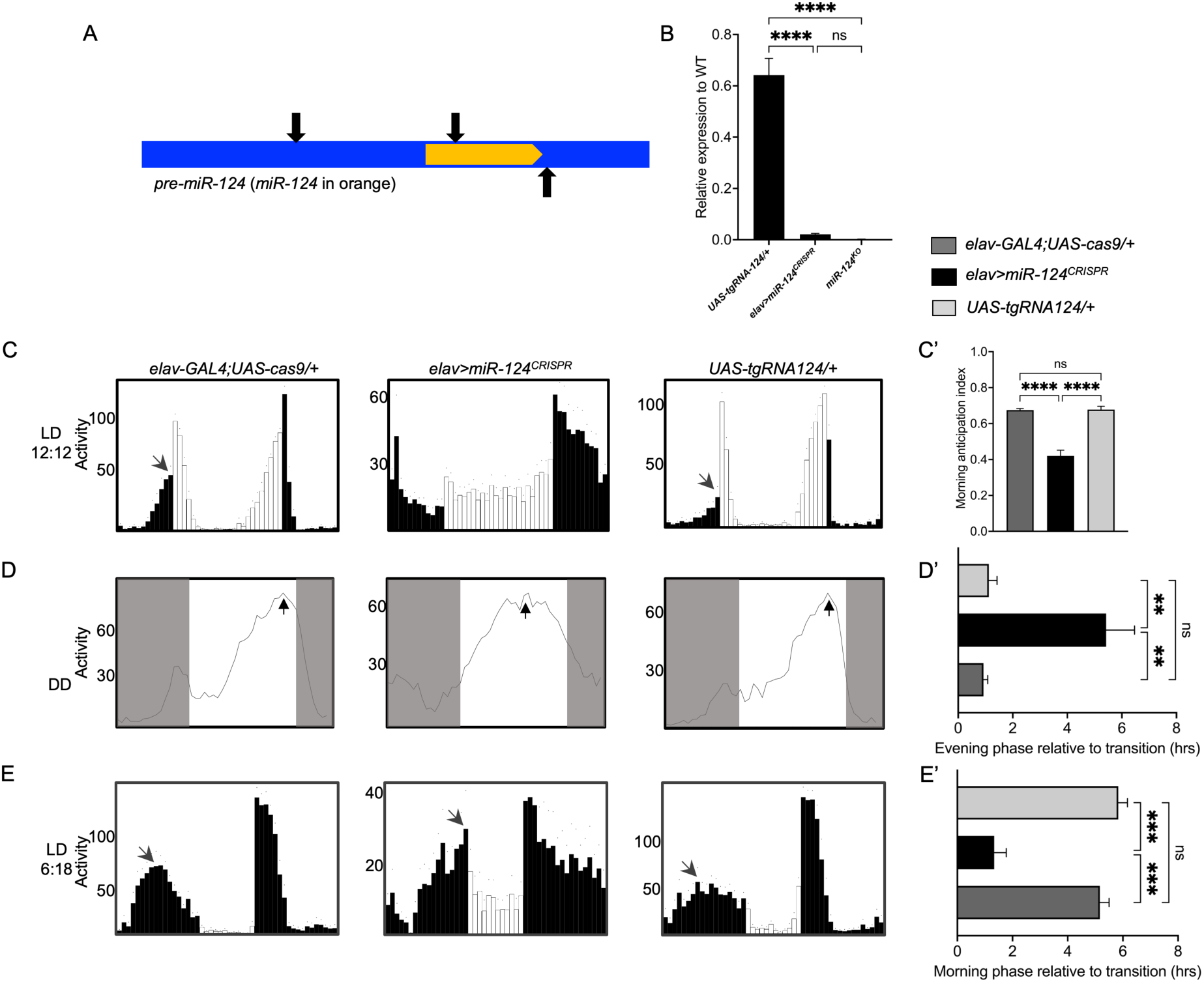
Pan–neuronal CRISPR-mediated disruption of *miR-124* phenocopies *miR-124^KO^*. *A*, Diagram showing three sgRNA target sites for pre-*miR-124*. *B*, Quantitative real-time PCR analysis of *miR-124* abundance in deletion mutants and controls. *elav* indicates the *elav-GAL4* driver, *miR-124^CRISPR^* refers to the genotype *UAS-Cas9/+;UAS-tgRNA124/+*. *C-E*, Averaged locomotor behavior of flies with *elav-Gal4*-driven disruption of *miR-124* in 12:12 LD (*C*), DD (*D*) and 6:18 LD (*E*) (C-E, n ≥ 10 per genotype). *C’-E’*, Quantifications of morning anticipation index in 12:12 LD (*C’*), evening peak phase under DD (*D’*) and morning peak phase in short photoperiods (*E’*) (N = 3 independent behavioral experiments). The legends for *C’-E’* are above the panels. Error bars represent SEM. One-way ANOVA followed by a Tukey post-hoc test was performed. **, P<0.0021; ***, P<0.0002; ****, P<0.0001; ns, not significant.

Taken together, these data demonstrate that *miR-124* acts in neurons to regulate circadian phase, consistent with its role as a conserved neuron-specific microRNA. Additionally, CRISPR/Cas9-mediated *miR-124* deletion in neurons is highly efficient and can thus be used to study *miR-124* temporal and spatial requirements.

### Developmental *miR-124* expression is essential for properly phased adult circadian behavior

Next, we attempted to narrow down the specific neurons required for *miR-124*-mediated circadian phase regulation by conducting a neuronal GAL4 screen. We crossed flies carrying both *UAS-Cas9* and *UAS-miR-124-sgRNA* transgenes with various GAL4 lines, including those targeting different subsets of clock neurons^47^ and non-clock neurons related to circadian output pathways^48–51^, to disrupt *miR-124* in these neurons. However, none of these GAL4 lines recapitulated the behavioral defects observed in pan-neuronal *miR-124^CRISPR^*flies. The *elav-GAL4* line we used is an enhancer trap line that fully mirrors the expression pattern of endogenous *elav*^52^, with expression first detected in the embryonic nervous system at stage 12. The absence of phenotype with the other GAL4 lines we tested might thus be because these lines are expressed too late during the development of the relevant neurons. To test whether *miR-124* is indeed required early during development, we employed the temporal and regional gene expression targeting (TARGET) system^53^ to induce *miR-124* deletion at different developmental stages. Pan-neuronal disruption during development advanced evening phase in DD, delayed morning phase under short photoperiods, and disrupted morning anticipation and light-on startle response under 12:12h LD cycles, recapitulating the behaviors seen with constitutive *miR-124* deletion (Fig. 5A-5C and 5A’-5C’). In contrast, disruption confined to the adult stage had no effect on circadian phase, with behaviors comparable to genetic controls (Fig. S2). *miR-124* is therefore required during development to control the phase of adult circadian behavior.

**Fig. 5.**
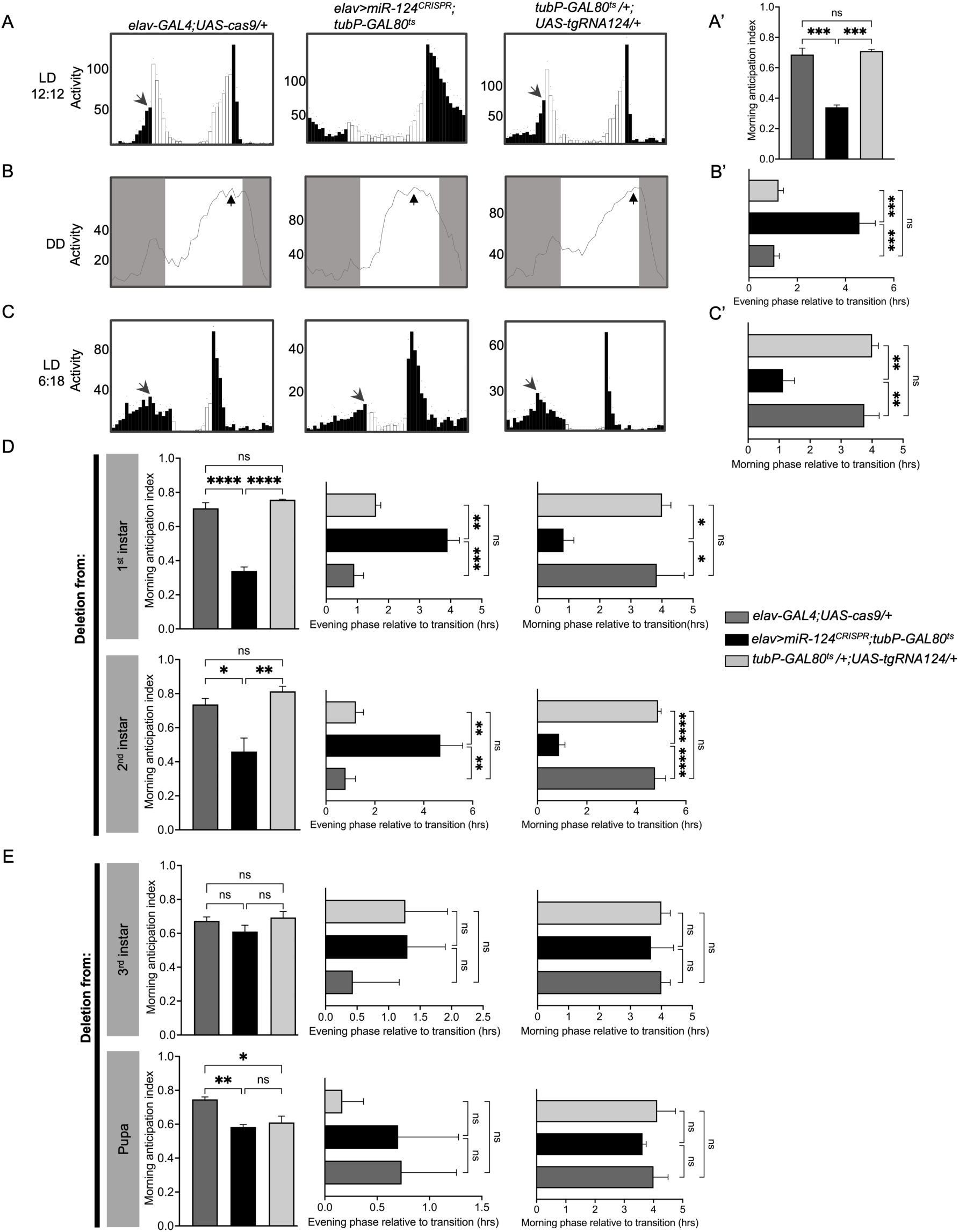
*miR-124* acts during development to regulate circadian phase. *A-C*, Averaged locomotor behavior of flies with embryonic *miR-124* deletion in 12:12 LD (*A*), DD (*B*) and 6:18 LD (*C*) (*A-B*, n ≥ 8 per genotype; *C*, n ≥ 9 per genotype). *A’-C’*, The quantification of morning anticipation index in 12:12 LD (*A’*), evening peak phase (hours) under constant darkness (*B’*) and morning peak phase (hours) in 6:18 LD (*C’*). *D*, Quantification of morning anticipation index (left), evening phase (middle) and morning phase (right) under 12:12 LD, DD and 6:18 LD for control flies and flies with 1^st^ instar larval and 2^nd^ instar larval deletion of *miR-124*. *E*, Quantification of morning anticipation index (left), evening phase (middle) and morning phase (right) under 12:12 LD, DD and 6:18 LD for control flies and flies with 3^rd^ instar larval and pupae stage deletion off *miR-124*. The legends for *A’-C’* and *D-E* are on the right side of panel D. See also Figure S2-S4 for additional data. Quantifications are means ± SEM in *A’-C’* and *D-E* (N = 3 independent behavioral experiments). One-way ANOVA with Tukey’s post hoc test. *, P<0.0332; **, P<0.0021; ***, P<0.0002; ****, P<0.0001; ns, not significant.

To determine more precisely when *miR-124* is required, we induced *miR-124* disruption at different developmental stages. Pan-neuronal *miR-124* deletion induced during the first or second instar larvae resulted in the same circadian phase defects as those observed with constitutive loss of *miR-124* (Fig. 5D and Fig. S3A-3B). In contrast, *miR-124* disruption induced during the third instar larvae or at the pupal stages had no effects on circadian behavior (Fig. 5E and Fig. S4A-4B). Thus *miR-124* is required around the second instar larval stage to determine adult behavior phase.

### Loss of *miR-124* leads to defective circadian network wiring

Given that *miR-124* functions during larval development and alters the activity of circadian neurons in *miR-124* mutants, we hypothesized that the loss of *miR-124* affects the wiring of the circadian neural network. To test this, we employed the unbiased anterograde trans-synaptic tracing system, *trans*-Tango, to map synaptically connected neurons within circadian circuits^54^.

Due to the reverse correlation between the postsynaptic trans-Tango signal and temperature over the 15–25°C range^54^, we raised and entrained our flies at 25°C to stringently assess neural connectivity. We first traced and defined circuits downstream of the s-LNvs by co-expressing the *trans*-Tango ligand and myrGFP using *pdf-GAL4*. In both WT and *miR-124^KO^*brains, we observed the typical innervation of the posterior dorsal protocerebrum by GFP-expressing s-LNvs, along with optic lobe and contralateral projections from l-LNvs (Fig. 6A and Fig. S5). However, in *miR-124^KO^*brains, the dorsal projections of the s-LNvs appeared to be less fasciculated, with projection branching out of the main bundle more frequently ventrally (Fig. S5B). Similar observations had been made previously^33^, although projection defects seemed more pronounced in the present study, perhaps because of improved imaging. Projection abnormalities might be the result of axon guidance defects, which were clearly observed in larvae. In WT larval brains, all s-LNv axons projected medially and terminate in the anterior region of the central brain (Fig. S6A). However, in *miR-124^KO^* brains, some s-LNv axons displayed aberrant wandering near the somas in the posterior brain region and even formed loop-like structures (Fig. S6B). Interestingly, these axon guidance defects were usually more pronounced in one hemisphere compared to the other (Fig. S6C). *miR-124* thus plays a role in s-LNv axon guidance.

**Fig. 6.**
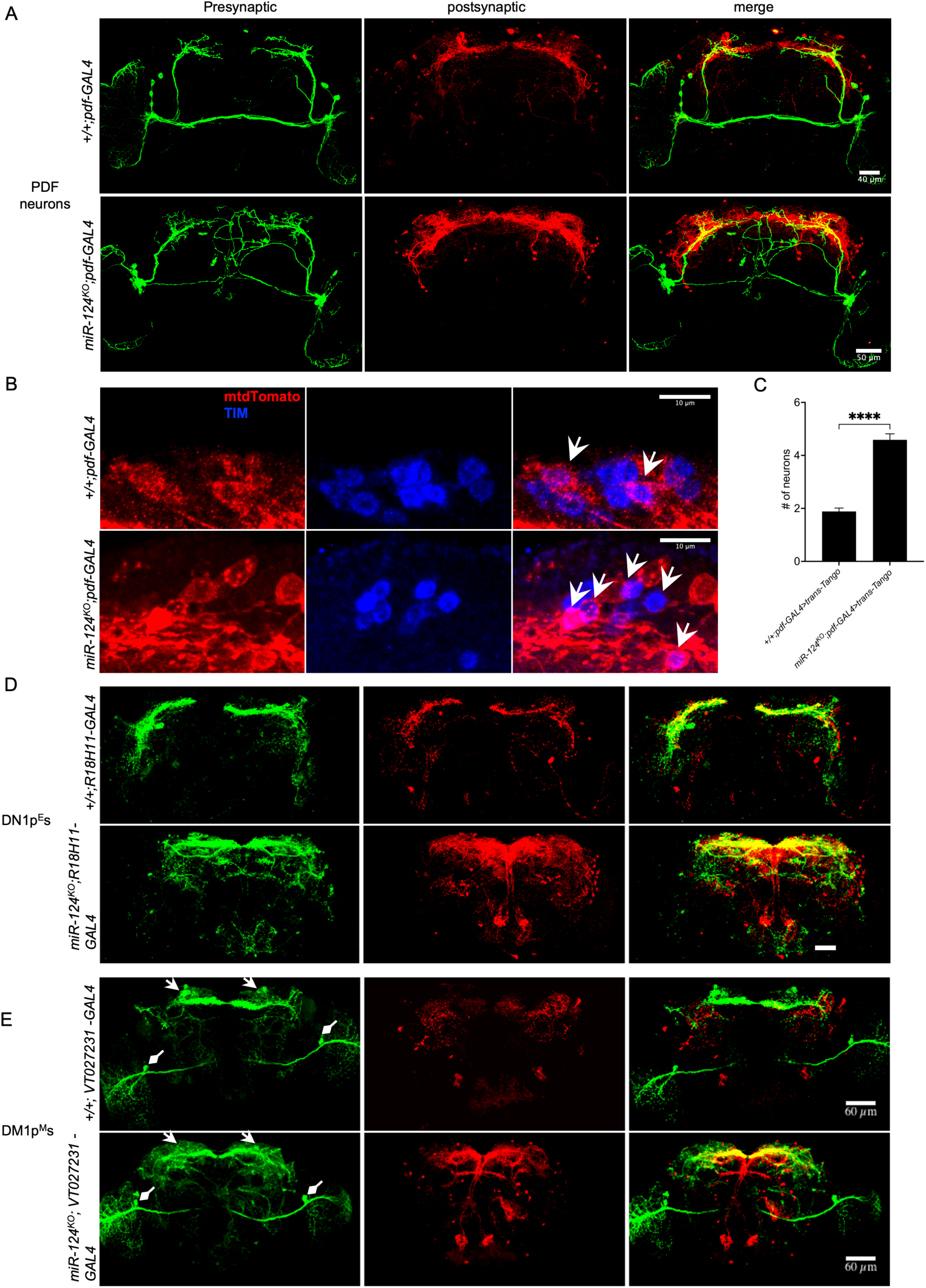
Loss of *miR-124* causes excessive synaptic connections in the circadian network. *A*, Postsynaptic partners of s-LNvs in WT and *miR-124^KO^* mutants revealed with *trans*-Tango. From left to right: Presynaptic s-LNvs (anti-GFP, green), postsynaptic *trans*-Tango signal (anti-HA, red), and merged images. Genotypes are indicated on the left side of the panels. The scale bars represent 40 μm. See also Figure S5-S6 for additional data. *B*, Direct synaptic contacts between s-LNvs and DN1ps in *A* revealed by co-staining with anti-TIM (blue). Brains were dissected at ZT21. White open arrowheads indicate the DN1ps directly targeted by s-LNvs. See also Figure S7. *C*, Quantification of the numbers of DN1ps contacted by s-LNvs per hemisphere. *D-E*, Postsynaptic partners of DN1p^E^s (*D*, scale bar, 50um) and DN1p^M^s (*E*) in WT and *miR-124^KO^* mutants revealed with *trans*-Tango. White arrows represent DN1p^M^, white diamond arrows indicate non-DN1p^M^s in *E*. See also Figure S8-S10 for additional data.

In addition to projections defects, adult *miR-124^KO^*brains displayed significantly increased Tomato signal in neurons postsynaptic to the s-LNvs relative to WT controls, suggesting that s-LNvs target a greater number of neurons in *miR-124^KO^* flies (Fig. 6A and Fig. S5). This increased signal was observed in both the dorsal part of the brain, and near the midline. Since the DN1ps are downstream of the s-LNvs and project fibers into the dorsal protocerebrum, we investigated whether the loss of *miR-124* affected their connections. By confirming their identity as clock neuron through TIMELESS (TIM) protein staining, we observed that in WT, 1.9±0.1 (mean±SEM) DN1ps per hemisphere were postsynaptic to the s-LNvs. However, in *miR-124^KO^*brains, 4.6± 0.2 DN1ps per hemisphere were targeted by the s-LNvs (Fig. 6B-6C and Fig. S7). This indicates that the loss of *miR-124* leads to increased synaptic connectivity between the s-LNvs and DN1ps.

Next, we sought to trace postsynaptic partners of both DN1p subtypes in wild-type and mutant flies. First, we co-expressed the *trans*-Tango ligand and myrGFP with the *R18H11-GAL4*, which specifically labels DN1p^E^s. In WT brains, most projections from *R18H11*-positive DN1ps are located in the dorsal brain, innervating the anterior optic tubercle (AOTU) and the pars intercerebralis (PI)^55,56^ (Fig. 6D and Fig. S8A).

However, in *miR-124^KO^* brains, in addition to these dorsal projections, numerous projections terminated more ventrally, near the antennal lobe (AL), and there were even sparse projections in the suboesophageal ganglion (SOG) (Fig. 6D and Fig. S8B). This extensive branching of DN1p^E^s in *miR-124^KO^* brains correlated with much broader and denser postsynaptic mtdTomato labeling. In WT brains, mtdTomato-positive projections were mainly restricted to the lateral and dorsal protocerebrum, whereas in *miR-124^KO^* brains, robust mtdTomato-positive projections were found throughout a large area of the central brain (Fig. 6D and Fig. S8). As a result, it was challenging to accurately identify specific synaptic connections. However, using TIM staining, we observed that some DN1p^E^s form synapses with each other. The loss of *miR-124* did not appear to affect this connectivity, as 4 out of 7-8 *R18H11-GAL4*-labelled DN1p neurons^57^ maintained contact with each other in both WT and *miR-124^KO^*flies (Fig. S9).

We then turned our attention to the DN1p^M^s. *VT027231-Gal4* labels the DN1ps which are sufficient to drive morning anticipation^18^, so we used this driver to express the *trans*-Tango ligand and myrGFP in the DN1p^M^s. Note that this driver is also expressed in a few non-DN1ps. In WT brains, GFP-expressing fibers from the DN1ps are primarily concentrated in the lateral and dorsal brain. However, in *miR-124^KO^* brains, these fibers spread extensively throughout the entire central brain (Fig. 6E and Fig. S10). This widespread and dense arborization corresponded with highly dense trans-Tango-dependent signals in the central brain of *miR-124^KO^* mutants. Although other clock neurons and non-clock neurons targeted by *VT027231-Gal4* may contribute to the widespread mtdTomato labeling, particularly in the ventral brain, the intense mtdTomato labeling around the DN1p^M^s strongly suggests that the loss of *miR-124* causes DN1p^M^s to form additional synaptic connections. Thus, both DN1p^E^s and DN1p^M^s are similarly affected by the loss of *miR-124*, exhibiting broader and denser projections in *miR-124^KO^* brains.

Collectively, these results suggest that the absence of *miR-124* leads to developmental miswiring within circadian circuits, likely disrupting neuronal activities in the circadian network and causing shifts in circadian phase.

## Discussion

*miR-124* is an evolutionarily conserved neuronal microRNA^58,59^. Its abundant expression in the CNS suggests an indispensable role in promoting and maintaining normal CNS function. In vertebrates, *miR-124* is critical for various physiological processes, including neural stem cell proliferation, neuronal differentiation, neurite growth and neuron migration^60,61^. Abnormal expression of *miR-124* or its target genes has been linked to numerous neurodevelopmental and neurodegenerative disorders^62,63^. Interestingly, *miR-124* appears to be less critical for the central nervous system of invertebrates. In *C. elegans*, *miR-124* knockout mutants are viable and show no gross abnormalities, nor are there evident defects in the number or differentiation of sensory neurons, where *miR-124* is predominantly expressed^64^. In *Drosophila*, previous work indicated that loss of *miR-124* leads only to mild neurodevelopmental defects. Specifically, the absence of *miR-124* leads to a reduction in the branch length and bouton numbers of neuromuscular junctions (NMJ) 6/7, as well as a decrease in the branch numbers of dendritic arborization (DA) neuron ddaE in the third larvae^65^. Additionally, it increases the variability in dendrite numbers of sensory neurons, particularly ddaD neurons, in larvae^66^. Despite these subtle developmental abnormalities, mutant flies display a range of physiological and behavioral impairments, including deficits in locomotion, flight ability, reduced lifespan, and decreased fertility^65–68^. Here, we report that *miR-124* plays a crucial role in the proper development and neuronal wiring of the *Drosophila* circadian neural network. Its absence leads to excessive synaptic connections within the circadian neural circuit, and between circadian and non-clock neurons. Since previously reported neural defects, observed in the larvae, were quite subtle, we were surprised by the extend of the wiring defects in the adult circadian neural network. Synaptic defects related to *miR-124* loss have been observed in other species. In *Aplysia californica*, *miR-124* modulates synaptic plasticity through CREB^69^, while in rat hippocampal neurons, *miR-124* controls input-specific synaptic plasticity via synaptopodin (SP)^70^. In mice, *miR-124* regulates dendritic morphogenesis and increases spine density in the olfactory bulb, highlighting its role in synaptic formation and plasticity^71^. Collectively, these data suggest that regulating synapse biology is an ancient and conserved function of *miR-124* across the animal kingdom.

The precise formation of synaptic connections between neurons is critical for proper information perception, processing, and transmission, which underpins all central nervous system functions^72^. Despite their distinct roles, the various clock neuron groups in the *Drosophila* brain are highly interconnected, allowing the flies to adapt to ever-changing environmental conditions^17,18,73^. The s-LNvs, the master pacemakers in *Drosophila*, maintain free-running rhythms under DD and are essential for anticipating dawn under LD conditions^15,16,74^. They also modulate the timing of the evening peak of locomotor activity^75^. Meanwhile, the DN1ps consist of subgroups that promote either morning or evening activity^18^. We focused on the neuronal connectivity of these two clock neuron subgroups, as the loss of *miR-124* affected both morning and evening phases in *miR-124^KO^* flies (Fig. 1). Our trans-Tango analysis revealed that in WT flies, only two DN1ps are postsynaptic to the s-LNvs (Fig. 6B-6C), which aligns with findings from the electron microscopy (EM) hemibrain connectome^76^, where s-LNvs are seen to form very few synaptic connections with other clock neuron groups^77^. This is consistent with previous studies showing that s-LNvs primarily communicate with DN1ps and other circadian groups through volumetric neuropeptide PDF secretion to set their pace^73,78,79^. In WT flies, the s-LNvs differentially activate the DN1p^M^s and inhibit the DN1p^E^s through PDF signaling^18^. Interestingly, the s-LNvs apparently use glycine, which usually inhibits neuronal activity, as its sole fast synaptic neurotransmitter^80^. In *miR-124^KO^*brains, we observed that the s-LNvs connect to 2-3 additional DN1ps. These additional synaptic connections may result in glycine signaling from s-LNvs overriding PDF signaling, thus inhibiting both DN1p^M^s and DN1p^E^s, since these neurons are connected to each other through electric synapses^81^.

Interestingly, the phase of s-LNv activity was altered in *miR-124* knockout animals, with activity shifted toward the middle of the day, rather than at the end of the night. This could be the result of altered neural input onto the s-LNvs, or *miR-124* could modulate the expression of circadian output genes that determine neural activity rhythms. It is noteworthy that although the loss of *miR-124* results in an increase in synaptic connections made by the s-LNvs and the DN1ps and drastically alters their activity pattern, only the phase of circadian rhythms is affected, not its period, nor the oscillations of neuronal molecular pacemakers^32,33^. For the s-LNvs, this fits with the observation that their neural activity can be eliminated without major consequences on the oscillation and phase of their circadian pacemaker^38^. However, in the case of *miR-124* mutants, the s-LNvs are rhythmically active, with an abnormal phase. Thus, the circadian molecular pacemaker of the s-LNvs is quite resistant to (or protected from) changes in neural activity, both in terms of period and phase. Since the s-LNvs function as the main circadian pacemaker neurons, this insulation might be needed to avoid overreaction to environmental inputs such as temperature changes^82^.

The DN1ps are controlled by the s-LNvs through PDF signaling, which maintain their rhythmicity and clock speed^18,41,83^. Altering the molecular clock in either s-LNvs or DN1ps readily shifts behavioral phase under DD, whereas similar molecular manipulations in LNds do not^18^, emphasizing the pivotal role of the s-LNv-DN1p axis in circadian phase regulation. The simultaneous re-activation of both morning and evening DN1ps significantly rescued circadian behavior in *miR-124^KO^*mutants (Fig. 3E-3G), indicating that neuronal activity and likely communication within DN1ps are essential for maintaining normal behavioral phases. Indeed, we (Fig. S9) and others observed interconnections between DN1ps^81^. Interestingly, in *miR-124* mutants, DN1ps activity is very low in both the E and M subtypes, and behavior phase correlated quite closely with s-LNv activity under DD: both showed a broad peak in the middle of the subjective day. This suggests that the s-LNvs can bypass the DN1ps to control behavior. The DN1ps respond to both light and temperature inputs^40^, and might thus be at time acutely inhibited. This could allow the s-LNvs to control behavior directly under specific environmental conditions.

Synapse formation requires a complex set of genes, including genes encoding cell-surface proteins, which differ among neuron types to specify their unique connectivity^84^. *miR-124* is predicted to directly regulates hundreds of genes, and the loss of *miR-124* affects the expression of thousands, particularly those involved in central nervous system (CNS) functions^66^. It is therefore likely that the wiring defects we observed is the result of the combined misregulation of multiple genes. Supporting this idea, reduction in BMP signaling - which impact multiple aspects of *Drosophila* development including synapse formation and function^85–89^-only partially rescued the evening *miR-14^KO^* phase phenotype, and this only under DD^32^. Clearly, *miR-124* is required quite early during development. Conditional rescue indicates that *miR-124* is required during mid or early larval development, and larval s-LNvs show frequent axonal guidance defects when *miR-124* is entirely missing. It is however curious that in adults, these striking defects have been largely corrected, with at best a minor increase in aberrant axonal projections (Fig. 6A and S5)^33^. It is therefore likely that the aberrant projections observed in larvae are pruned during pupation, or perhaps even earlier since we usually observed axonal guidance defects in only one of two brain hemispheres. The main defect of adult s-LNvs thus appear to be the excess connectivity with DN1ps and other unidentified neurons. The origin of this hyper-connectivity, also present in DN1ps, is presently unclear. It could be the result of more subtle axon guidance defects than those observed at the larval stages, or insufficient pruning.

Our findings demonstrate that *miR-124* regulates circadian phase of rhythmic behavior by coordinating the development of circadian neural circuits. They reveal the importance of proper circuit wiring in the control of behavior phase. Defective neural connectivity may also help explain other physiological and behavioral abnormalities caused by the loss of *miR-124*, not only in *Drosophila* but potentially also in other animals, given that *miR-124*’s mature sequence and brain-enriched expression are conserved from worms to humans.

### Experimental Procedures Drosophila strains and husbandry

All flies were reared on standard low yeast brown food at 25°C under 12:12 light/dark cycles unless otherwise specified. The following fly lines were used for this study: *w1118*; *miR-124^KO^* and rescued *miR-124*^KO^ (*39N16*) were generated by Sun et al.^66^, *y w; Pdf-GAL4/CyO* was described by Renn et al.^74^, *UAS-Kir2.1* was a gift from Dr. Michael Rosbash, *UAS-dTrpA1* was from Dr. Paul Garrity. *Clk4.1M-GAL4 was* generated by Zhang et al^40^. *UAS-uMCas9* (VDRC 340009) and *VT027231-GAL4* (VDRC 205530) were obtained from the Vienna *Drosophila* Resource Center (VDRC). The following strains were ordered from the Bloomington Stock Center*: Clk856-GAL4* (BDSC 93198), *R18H11-GAL4* (BDSC 48832), *UAS-CaMPARI2* (BDSC 78316), *elav-GAL4;tubP-GAL80^ts^*(BDSC 67058), *UAS-cas9* (BDSC 58985), *UAS-myrGFP.QUAS-mtdTomato-3xHA;trans-Tango* (BDSC 77124), *w[*];trans-Tango* (BDSC 77123), *UAS-myrGFP P,QUAS-mtomato-3xHA* (BDSC 77479), *elav-GAL4;UAS-cas9.P2* (BDSC 67073).

### Fly line generation

We constructed the *UAS-miR-124-sgRNA* plasmid by following the protocol published by F. Port and S. L. Bullock^45^. Briefly, three gRNAs targeting *pre-miR-124* were selected according to flyCRISPR (https://flycrispr.org/target-finder/). GAL4/UAS controlled gRNA-expressing pCFD6 vector (Addgene #73915) was digested with BbsI-HF (NEB #R3539S) and gel purified. Two inserts carrying the three gRNAs were generated by PCR reactions using pCFD6 as the template. The resulting two PCR fragments and linearized pCFD6 were then assembled in a single reaction by NEBuilder HiFi DNA Assembly (NEB #E2621S). The construct was inserted into the third chromosome using phiC31 integrase (attP2, BL8622) by the BestGene Inc (Chino Hills, CA).

The following gRNA and primers sequences were used:

gRNA1: TTTCTCCTGGTATCCACTGTAGG

gRNA2: TATTTCCACCATAAGGCACGCGG

gRNA3: TGTAGAACTGCGTTCGCTCTTGG

gRNA1F: CGGCCCGGGTTCGATTCCCGGCCGATGCATTTCTCCTGGTATCCACTGT GTTTCAGAGCTATGCTGGAAAC

gRNA1R: CGTGCCTTATGGTGGAAATA TGCACCAGCCGGGAATCGAACC

gRNA2F: TATTTCCACCATAAGGCACGGTTTCAGAGCTATGCTGGAAAC

gRNA2R:ATTTTAACTTGCTATTTCTAGCTCTAAAACAGAGCGAACGCAGTTCTACATGCACCAGCCGGGAATCGAACC

### RNA extraction and qRT-PCR

*miR-124* expression was measured with the miScript PCR System (Qiagen). Small RNA including miRNA from fly heads was extracted using miRNeasy Micro Kit (QIAGEN, Cat No. 217084) following the manufacturer’s instructions. The cDNA was synthesized using miScript II RT kit (QIAGEN, Cat No. 218161) according to the supplier’s protocol. To quantify mature *miR-124*, qRT-PCR was performed by using miScript SYBR Green PCR Kit (QIAGEN, Cat No. 218073) and Bio-Rad thermocycler CFX96. Reaction mix contained a universal reverse primer and a *miR-124*-specific forward primer provided by QIAGEN.

### Behavior recording and data analysis

Individual adult male flies (aged 2-7 days) were used to measure locomotor activity rhythms. In most experiments, flies were entrained to a 12:12 LD cycle for 3 days followed by at least 6 days of DD at 25**°**C, in Percival I36-LL incubators. To test flies under long or short photoperiod, 6h:18h LD or 16h:8h LD cycles at 25**°**C were used. Locomotor activity was recorded in Trikinetics *Drosophila* Activity Monitors (Waltham, MA). Group activity for each genotype was analyzed using the FaasX software (<u>Grima et</u> <u>al., 2002</u>). Under LD conditions, data from three days were analyzed. For DD conditions, data from 4-5 days of activity were used to determine the evening activity phase. Morning anticipation index was determined with the dedicated FaasX routine, which divides the average activity 3h prior to lights-on with the activity 6h before lights-on.

Under 6:18 LD cycles, the phase of the M-peak was quantified as the highest average activity time point before ZT0. Similarly, the E-peak phase was calculated as the highest average activity time point before CT12. All phase quantifications were manually performed by an observer who was blinded to the genotypes, using FAAS graphic outputs.

The Morning/Evening (M/E) activity ratio (figure 3F”) was calculated by dividing the average morning activity (CT0-2) with the average activity over 2.5 hours centered around the evening peak of activity.

### Conditional *miR-124* knock-out using the TARGET system

To induce *miR-124* knock-outs in adult flies only with the TAGRET system, flies of experimental genotypes were maintained at 18°C (restrictive temperature, GAL80ts active, no gRNA expression) during their entire development, and moved to 29°C (permissive temperature, GAL80ts inactive, gRNA expression) three days prior to the beginning of the locomotor activity assays. Control flies were exposed to the same temperature conditions. To induce the knock-out early during development, flies were exposed to 29°C throughout their development and adulthood.

To induce *miR-124* deletion at specific developmental stages, parental flies were kept at 25°C for 24 hours to allow for mating and egg laying. Eggs were then moved to 18°C to activate GAL80^ts^ repression and thus block expression of the *miR-124* gRNAs. At specific developmental stages, animals were transferred to 29°C until adulthood, to inactive GAL80ts and thus induce gRNA expression. Developmental stages were visually assessed, and genetic controls were raised alongside experimental flies to account for any temperature effects on development.

### Immunohistochemistry and Image Analysis

Third instar larvae or adult fly heads were dissected in dissection buffer (KCl 13.6g, NaCl 2.7g, CaCl_2_ 0.33g, Tris 1.21g in 1L water, pH 7.2) and immediately fixed in 4% paraformaldehyde diluted in PBS for 30 minutes at room temperature. Following two quick rinses, brains were washed three times with PBST (0.1% Triton X-100) for 10 minutes each. The brains were then blocked in 10% normal donkey serum in PBST for one hour at room temperature, followed by incubation with the primary antibodies at 4°C overnight. Afterward, the brains were washed six times in PBST (20 minutes each) at room temperature and incubated with the secondary antibodies at 4°C overnight. Finally, the brains were washed six times in PBST (20 minutes each) at room temperature and mounted on glass slides using Vectashield mounting medium (Vector Laboratories).

Primary antibodies were: mouse anti-PDF (Developmental Studies Hybridoma Bank, PDF; 1:400), guinea pig anti-TIM^90^ (1:400), rabbit anti-GFP (Thermo Fisher Scientific, A11122; 1:1000), mouse anti-HA (Covance, MMS-101P; 1:250). Secondary antibodies were diluted at 1:200-400 and were as follows: donkey anti-rabbit Alexa Fluor® 488, Goat anti-mouse Rhodamine Red, donkey anti-mouse Alexa Fluor® 546, donkey anti-guinea pig Alexa Fluor® 647. For TIM staining, flies were dissected at ZT 21 after 3 days of LD entrainment.

All brains were imaged using a Zeiss LSM700 confocal microscope with ZEN software, or Zeiss LSM800 confocal microscope with ZEN blue software, keeping the laser settings constant within each experiment. Images were processed using Fiji software (http://fiji.sc). To count the number of DN1ps postsynaptically connected with the s-LNvs, an observer blind to genotypes counted the number of DN1s that showed postsynaptic trans-tango signal surrounding nuclear TIM signal.

### CaMPARI imaging

Flies expressing UAS-CaMPARI crossed with *clk856-GAL4*^91^ were used to monitor calcium dynamics in clock neurons. Male progenies were entrained in 12h:12h light-dark cycles at 25°C for 4–5 days before transitioning to constant darkness (DD) at the same temperature. Flies were collected at the indicated time points during the first DD.

Flies were mounted on Petri dishes and exposed to UV light (395 nm, 10mW/cm2) for 5 minutes to induce CaMPARI’s calcium-dependent photoconversion. Their brains were then immediately dissected in cold extracellular fly buffer^92^ under dim red light. Brain tissues were then imaged using a Zeiss 800 confocal microscope.

Analysis involved processing images in ImageJ, with background subtraction to measure the green (F_green) and red (F_red) fluorescence intensities in target neurons. The CaMPARI photoconversion efficiency was calculated as the F_red/F_green ratio.

### Data and material availability

Any additional information and materials reported in this paper are available from the corresponding authors upon request.

## Acknowledgments

We thank Dr. Eric Lai for the *miR-124*^KO^ and genomic rescue flies, and Dr. Maria de la Paz Fernandez for helpful discussions. We also thank Dr. Michael Rosbash, the Bloomington Drosophila Stock Center, and the Vienna Drosophila Resource Center for *Drosophila* stocks and reagents. This work was supported by MIRA award 1R35GM118087 from the National Institute of General Medicine Sciences (NIGMS) to P.E.

## Author Contributions

Conceptualization, Y.X. and P.E.; methodology, Y.X., C.C. and P.E.; investigation, Y.X and C.C.; writing – original draft, Y.X.; writing – review & editing, Y.X., C.C. and P.E., funding acquisition, P.E.; supervision, P.E.

## Declaration of generative AI and AI-assisted technologies in the writing process

During the preparation of this work the authors used ChatGPT in order to ensure grammatical accuracy and improve clarify. After using this tool/service, the authors reviewed and edited the content as needed and take full responsibility for the content of the publication.

## Figure Legends

**Fig. S1. Activity profile of *w1118*, *miR-124^KO^* and *39N16* under 18:6 LD cycle.**

Related to Figure 1E. Time of evening peak activity is indicated on the graph (n ≥ 16 per genotype), as number of hours before lights-off. Genotypes are indicated under the panels.

**Fig. S2. Adult deletion of *miR-124* does not affect circadian locomotor activity.**

Related to Figure 4-5. *A-C*, Averaged locomotor behavior of flies with *miR-124* deletion induced during the adult stage in 12:12 LD (*A*), DD (*B*) and 6:18 LD (*C*) (*A*, n ≥ 16 per genotype; *B*, n ≥ 12 per genotype; *C*, n ≥ 16 per genotype). *A’-C’*, Quantification of morning anticipation index in 12:12 LD (*A’*), evening peak phase (hours) under DD (*B’*) and morning peak phase (hours) in 6:18 LD (*C’*) (N = 3 independent behavioral experiments). The legends for *A’-C’* are under the panels. Error bars represent SEM. One-way ANOVA followed by a Tukey post-hoc test was performed. ns, not significant.

**Fig. S3. Representative locomotor activity profiles of flies with *miR-124* deletion induced during the 1^st^ and 2^nd^ instar larval stages.**

Related to Figure 5D. Representative locomotor activity profiles for control flies and flies with 1^st^ instar larval (*A*) and 2^nd^ instar larval (*B*) deletion of *miR-124* under 12:12 LD (upper), DD (middle) and 6:18 LD (bottom). n ≥ 8 per genotype.

**Fig. S4. Representative locomotor activity profiles of flies with *miR-124* deletion induced during the 3^rd^ instar larval and pupal stages.**

Related to Figure 5E. Representative locomotor activity profiles for control flies and flies with 3^rd^ instar larval (*A*) and pupal (*B*) deletion of *miR-124* under 12:12 LD (upper), DD (middle) and 6:18 LD (bottom). n ≥ 8 per genotype.

**Fig. S5. Connectivity of s-LNvs.**

Related to Figure 6A. *A-B*, Confocal images of five adult brains displaying the post-synaptic neurons (red, anti-HA) of s-LNvs (green, anti-GFP) in WT (*A*) and *miR-124^KO^*mutant brains (*B*).

**Fig. S6. s-LNvs axonal projection patterns in third instar larval brains.**

Related to Figure 6A. *A*, Projection patterns of s-LNvs neurons from six WT larval brains. *B-C*, Projection patterns of s-LNvs from six *miR-124^KO^* larval brains. s-LNvs from one hemisphere display severe axon guidance defects (*B*), the projection patterns from the other hemisphere of the same brains appear normal (*C*). The s-LNvs were visualized with PDF antibody staining (green). Open arrowheads indicate fibers that form loop-like structures.

**Fig. S7. Colocalization of trans-tango and TIM signal**

Partial z-stacks focused on the DN1p cluster to illustrate the co-localization of TIM (blue) and mtdTomato (red), related to Figure 6B. To highlight the intact structure of neurons, with mtdTomato signals surrounding TIM, a limited number of image slices were used to illustrate the co-localization of mtdTomato and TIM. *A*, Two DN1ps are post-synaptic to the s-LNvs in one hemisphere of a WT brain. *B*, Eight DN1ps are post-synaptic to the s-LNvs in one hemisphere of a *miR-124^KO^* mutant brain.

**Fig. S8. Synaptic connections of DN1p^E^s**

Related to Figure 6D. *A-B*, Confocal images of five adult brains showing post-synaptic neurons (red, anti-HA) of DN1p^E^s (green, anti-GFP) in WT (*A*) and *miR-124^KO^* mutant flies (*B*).

**Fig. S9. *miR-124* knock-out does not affect synaptic contacts between DN1p^E^s.**

Related to Figure 6D. *A*, Representative confocal images of direct synaptic contacts between DN1p^E^s revealed by trans-Tango. The brains were dissected at ZT21 and stained with anti-HA (red), anti-GFP (green) and anti-TIM (blue) to reveal that some DN1p^E^s contact each other. Genotypes are indicated on the left side of the panels. *B*, Average number of DN1p^E^s that connect with each other in each hemisphere. This number was always 4 in the 6 hemispheres that were imaged, hence the absence of error bars.

**Fig. S10. Synaptic connections of DN1p^M^s**

Related to Figure 6E. *A-B*, Confocal images of five adult brains showing the post-synaptic neurons (red, anti-HA) of DN1p^M^s (green, anti-GFP) in WT (*A*) and *miR-124^KO^*mutant flies (*B*). White arrows represent DN1p^M^s, white diamond arrows indicate non-DN1p^M^.

