## Supplementary figures and images for "*miR-124* acts during *Drosophila* development to determine the phase of adult circadian behavior"

### Supplemental figure S1

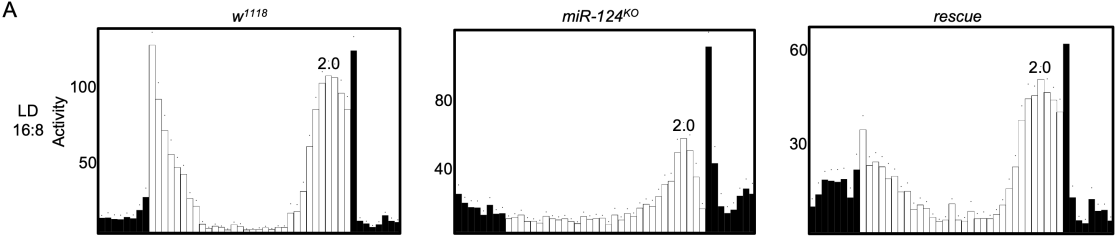

### Supplemental figure S2

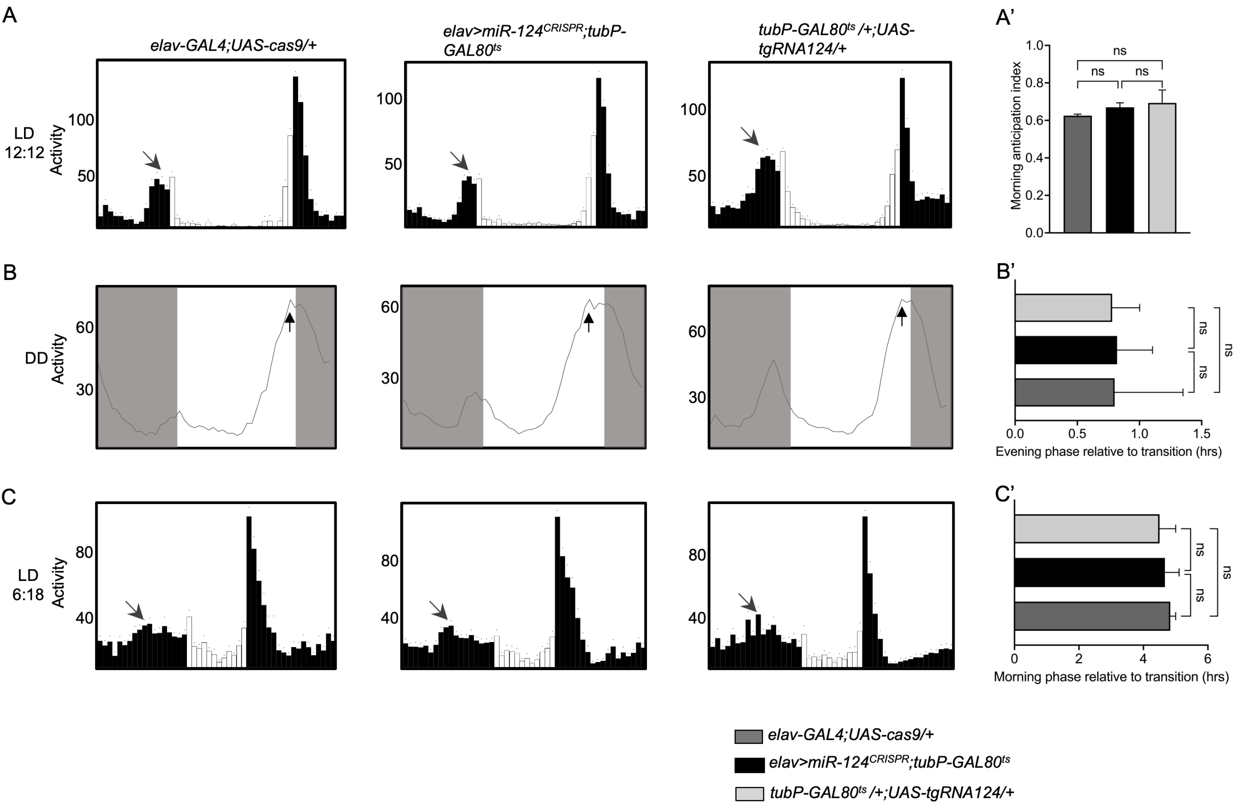

### Supplemental figure S3

Fig. S3

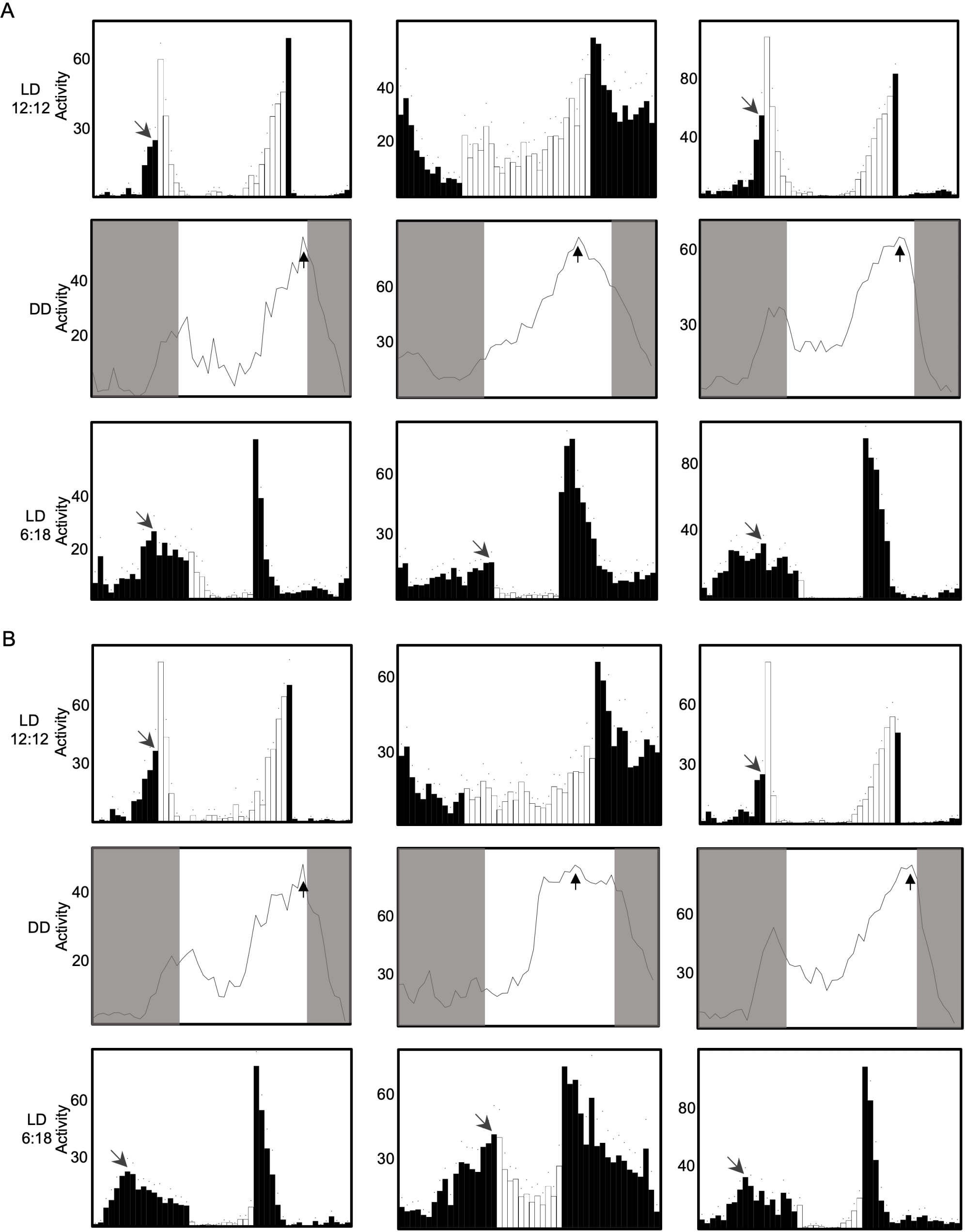

### Supplemental figure S4

Fig. S4

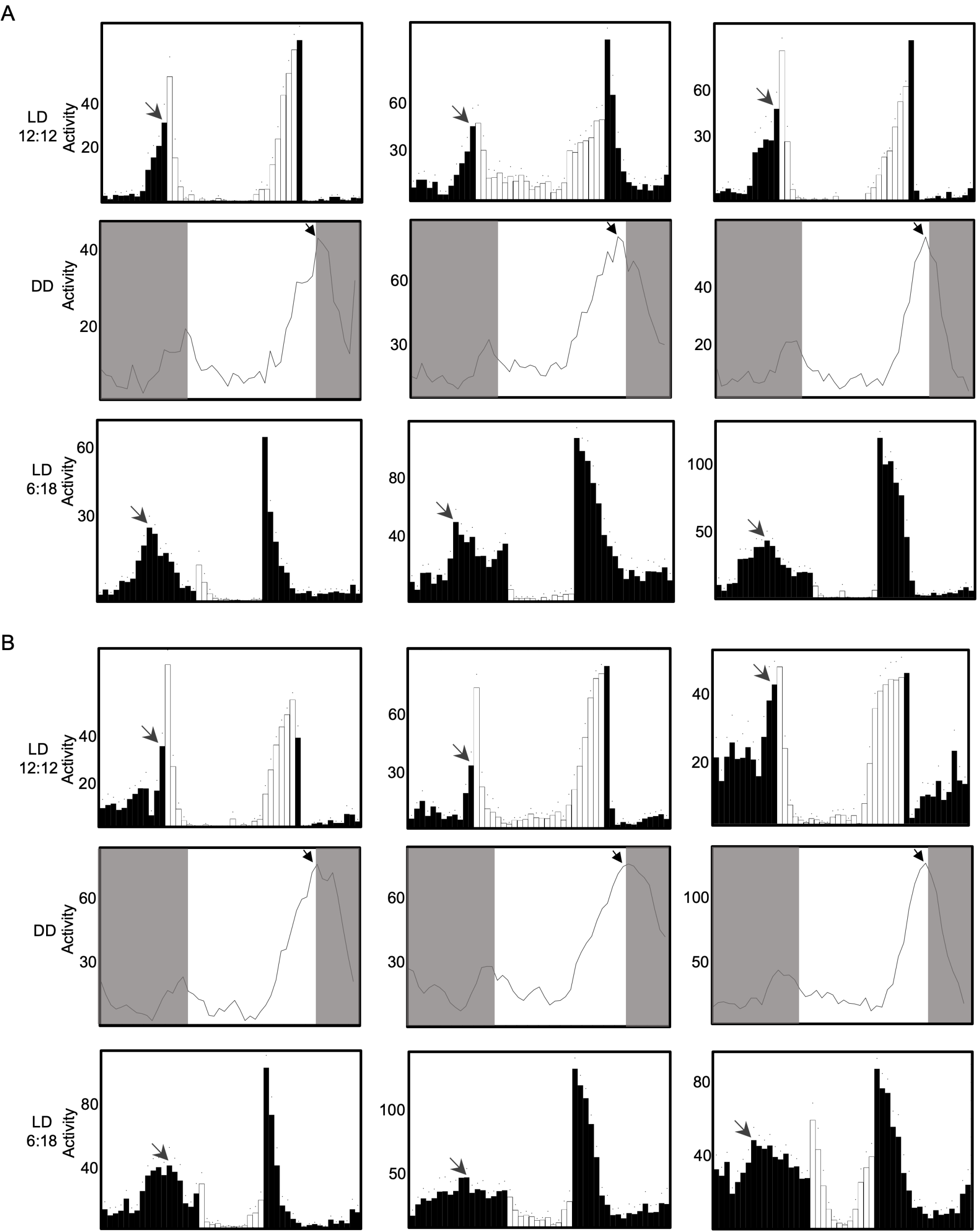

### Supplemental figure S5

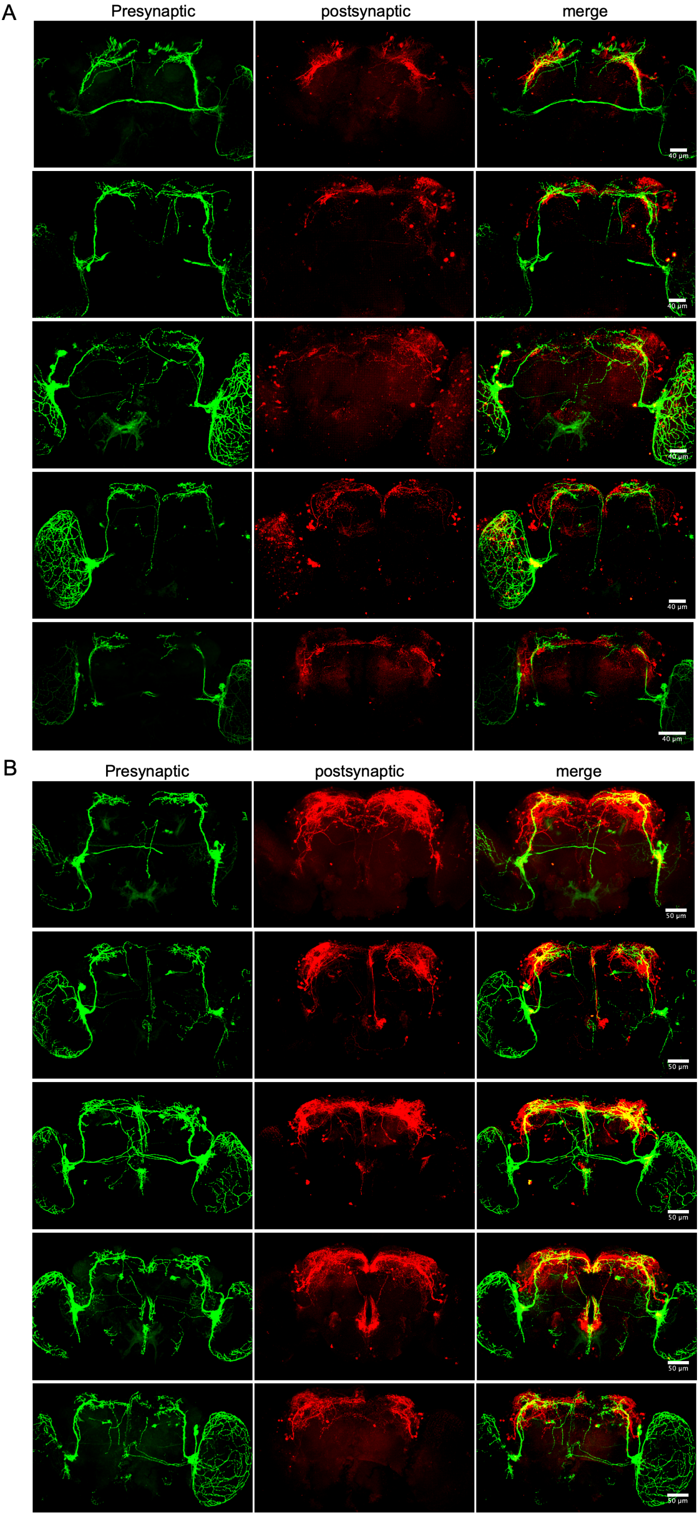

### Supplemental figure S6

Fig. S6

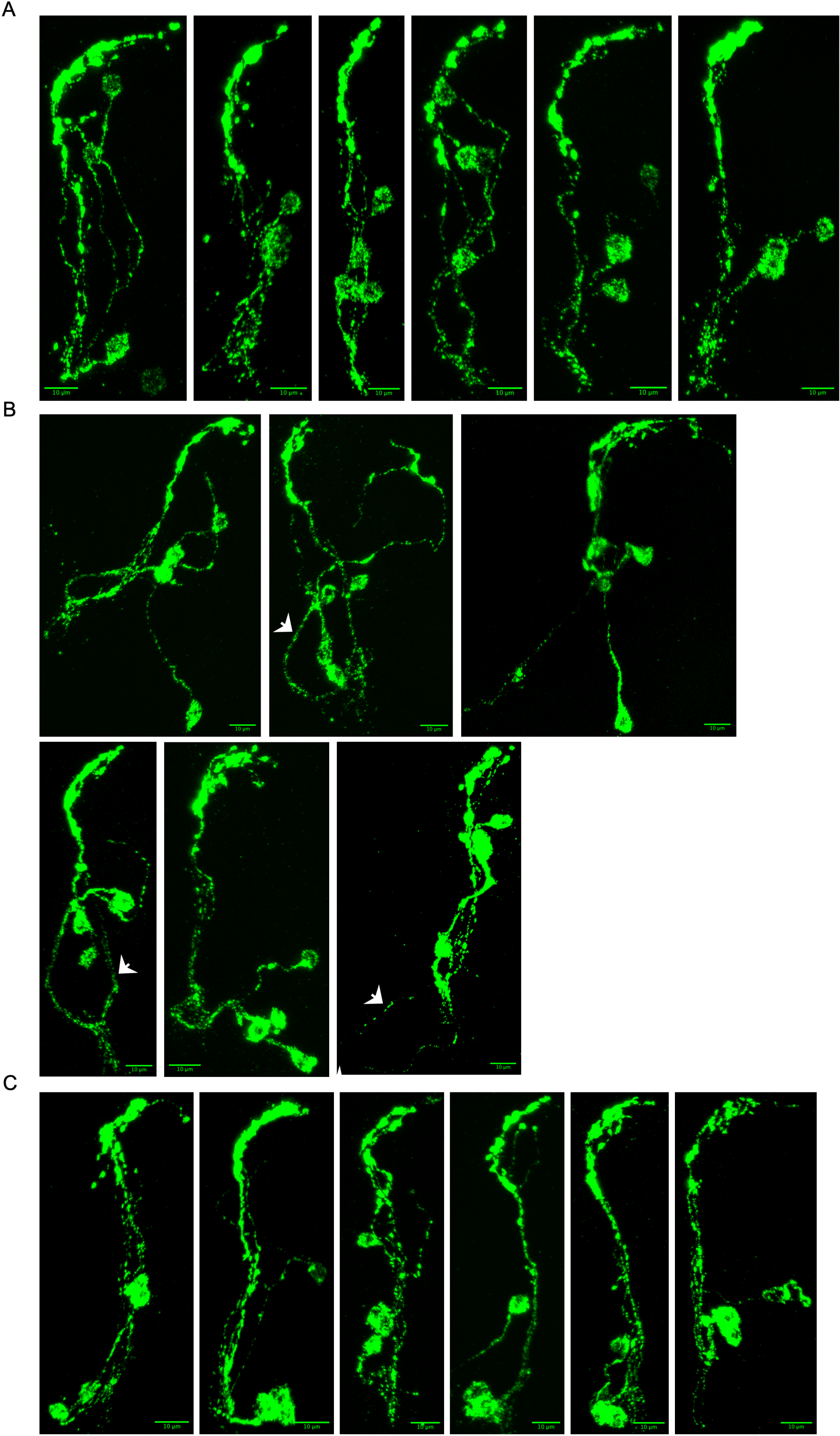

### Supplemental figure S7

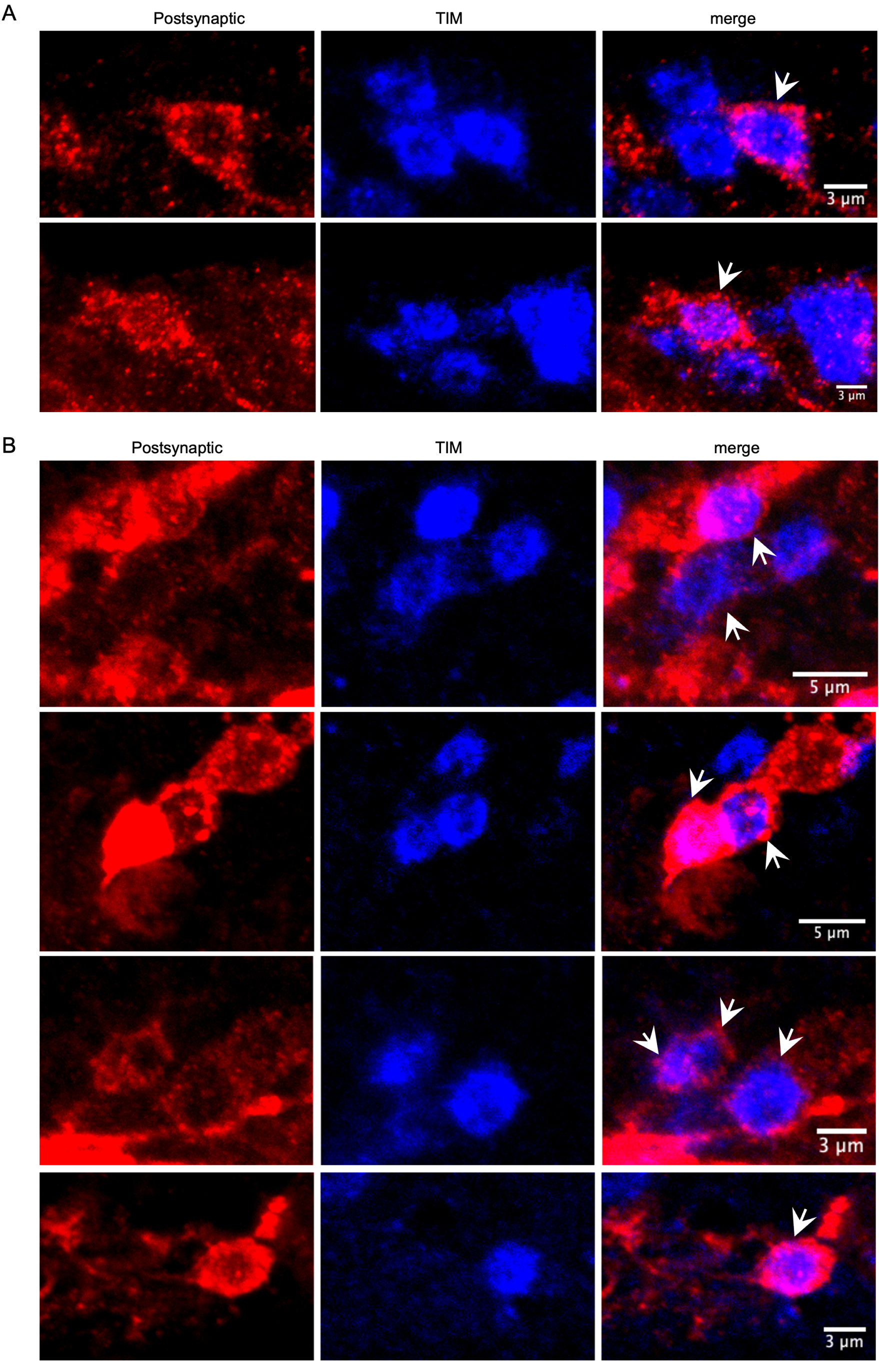

### Supplemental figure S8

Fig. S8

A

Presynaptic | postsynaptic | merge

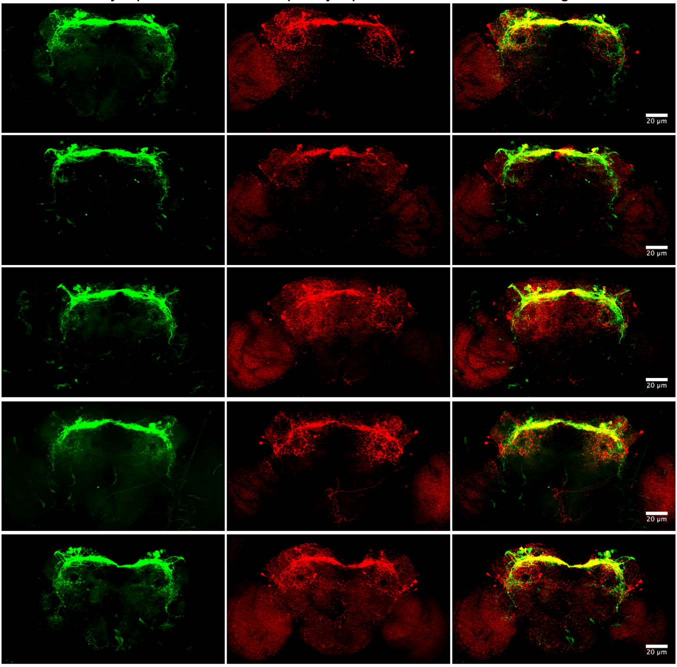


B

Presynaptic | postsynaptic | merge

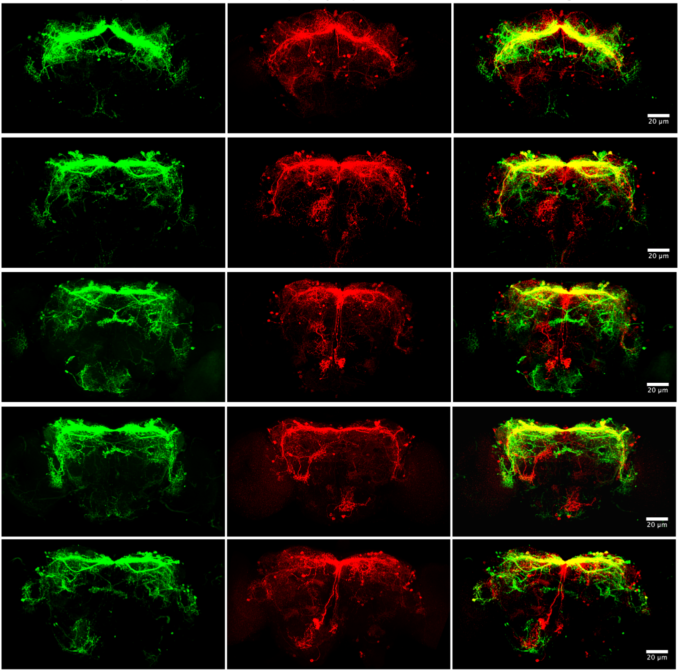

### Supplemental figure S9

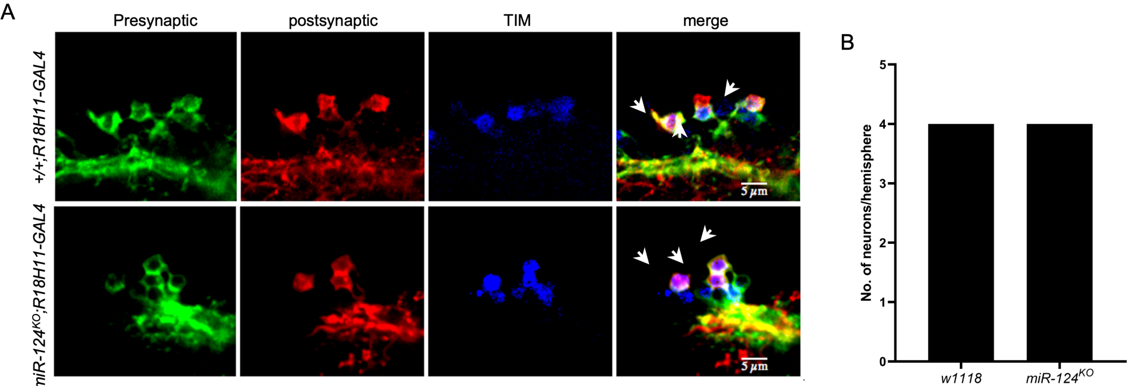

### Supplemental figure S10

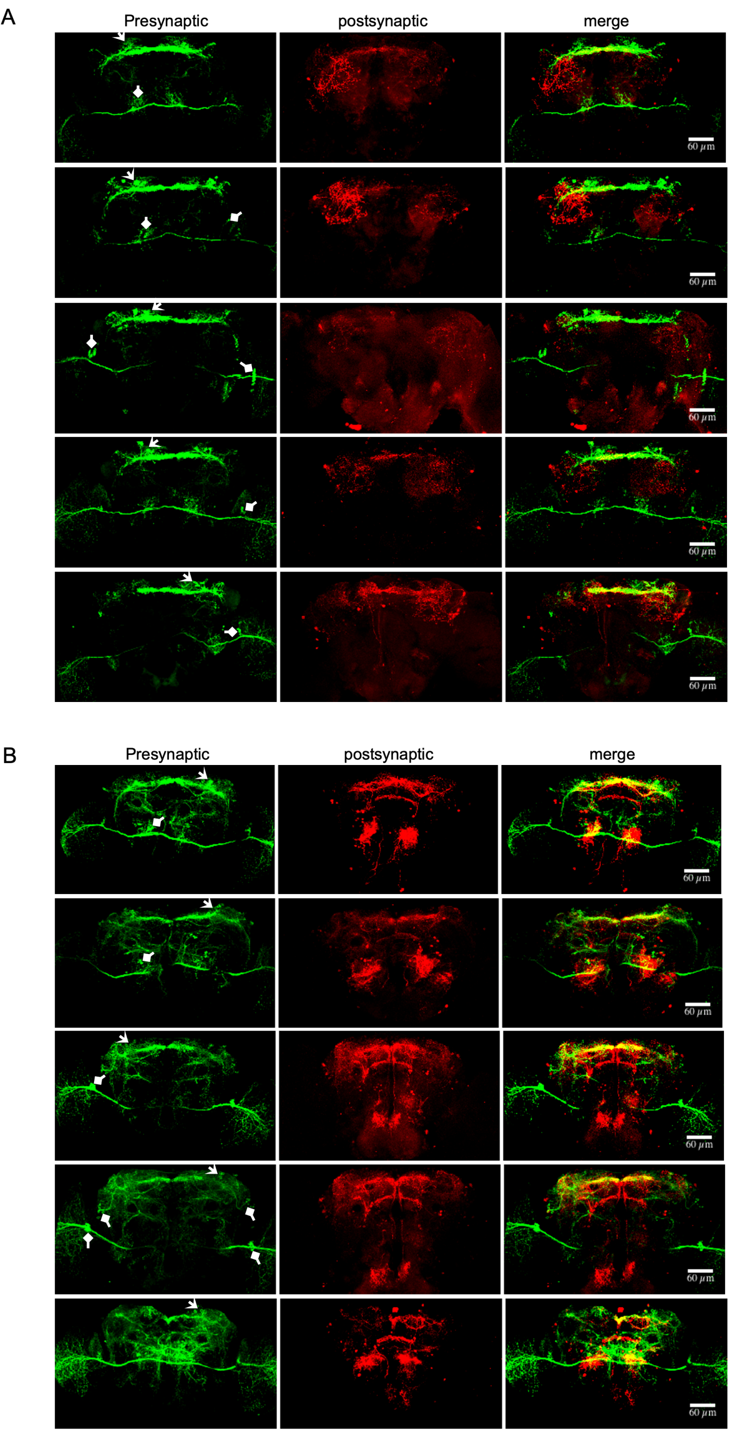
